# *norCBD* disruption affects the H2-type six secretion system and multiple virulence factors in *Pseudomonas aeruginosa*

**DOI:** 10.1101/2025.09.23.677791

**Authors:** Md Mahamudul Haque, Sara Badr, Kangmin Duan

## Abstract

The type six secretion system (T6SS) is a macromolecular weapon used by many Gram-negative bacteria. The T6SS functions as a needle injection system that delivers effector proteins directly into neighboring bacterial cells, thereby affecting their gene expression and physiological processes. *Pseudomonas aeruginosa* possesses three distinct T6SSs, designated as H1-, H2-, and H3-T6SS. Although extensive studies have been carried out on these T6SS systems in recent years, the regulatory mechanisms of T6SS remain incomplete. Here, we report the identification of *norCBD* as an operon that modulates the transcriptional activity of H2-T6SS. Both transposon insertion at *norCBD* and the deletion of the *norCBD* genes significantly reduced the CTX-*H2*-T6SS reporter activity. The *norCBD* operon encodes nitric oxide reductase (NorCBD), which reduces nitric oxide (NO) to nitrous oxide (N₂O), a crucial step in reducing the toxic levels of intracellular NO and facilitating anaerobic respiration. As the transcriptional regulatory Dnr activates H2-type VI secretion system (H2-T6SS) in response to NO, experiments were carried out to examine whether *norCBD* deletion caused intracellular NO accumulation, which in turn disrupted Dnr-dependent regulation of H2-T6SS and virulence factors. The NO levels and Dnr-regulated gene expression were measured, and several virulence-related phenotypes were examined. The effects of NO donor sodium nitroprusside (SNP) and NO scavenger carboxy-phenyl-tetramethylimidazolineoxyl (CPTIO) were also tested. The data obtained indicate that deletion of *norCBD* led to intracellular NO accumulation, reduced H2-T6SS expression, and affected motility, pyocyanin production, and biofilm formation. Complementation of *norCBD* on a plasmid in the deletion mutant was able to restore H2-T6SS expression and the examined phenotypes to the wild-type levels. Treatment with CPTIO also restored H2-T6SS expression in the PAO1(*ΔnorCBD*). These results indicate that NorCBD plays a critical role in maintaining NO homeostasis that is necessary for effective Dnr-mediated gene regulation and multiple virulence-related traits, highlighting the importance of redox balance in coordinating respiration and pathogenesis in *P. aeruginosa*.

## Introduction

*Pseudomonas aeruginosa* is a Gram-negative pathogen that plays a major role in chronic polymicrobial lung infections in cystic fibrosis (CF) patients (Nakatsuka, Matsumoto, Inohara, & Nunez, 2023). It is also a common cause of pneumonia, urinary tract infections, and burn wound infections. Antimicrobial-resistant *P. aeruginosa* has been recognized by the World Health Organization (WHO) as a critical priority pathogen that poses serious risks to global public health (Willyard, 2017).

The pathogenic success of *P. aeruginosa* is largely attributed to its arsenal of virulence factors (Qin et al., 2022). In particular, it employs specialized secretion systems to deliver toxins into host cells and competing microbes. Among these, the Type VI Secretion Systems (T6SSs) act as contractile nanomachines that mediate intercellular interactions by injecting toxic effectors into neighboring cells (Gallique, Bouteiller, & Merieau, 2017). T6SSs have been implicated not only in interbacterial competition but also in metal ion acquisition (Chen et al., 2016; Cianfanelli, Monlezun, & Coulthurst, 2016; Wang et al., 2015) and biofilm formation (Alteri et al., 2013).

*P. aeruginosa* possesses three functionally distinct T6SSs: H1-T6SS, H2-T6SS, and H3-T6SS. H1-T6SS is primarily involved in intraspecies competition and delivers at least seven identified toxins (Tse1– Tse7) into rival bacteria (George, Narayanan, Tejada-Arranz, Plack, & Basler, 2024; Hood et al., 2010). In contrast, H2-T6SS and H3-T6SS are capable of targeting both prokaryotic and eukaryotic cells (Lin et al., 2017). Despite their critical roles, the regulatory mechanisms that govern T6SS activity, particularly for H2- and H3-T6SS, remain incompletely understood.

To identify genes involved in H2-T6SS regulation, we employed transposon mutagenesis using a CTX-H2-T6SS reporter system. This screen revealed that disruption of the *norCBD* operon significantly decreased H2-T6SS reporter activity. The *norCBD* operon encodes the nitric oxide reductase complex (NorCBD), which catalyzes the reduction of toxic nitric oxide (NO) to nitrous oxide (N₂O), a key step in anaerobic respiration.

The NorCBD complex is a cytochrome bc-type nitric oxide reductase. NorB forms the membrane-integrated catalytic subunit, NorC functions as a cytochrome c electron carrier, and NorD is thought to facilitate proper assembly and stabilization of the enzyme complex (Gao et al., 2016). Expression of the *norCBD* operon is activated by the nitric oxide-responsive transcription factor Dnr, which is itself regulated by Anr, the master transcriptional regulator that responds to low oxygen tension (Arai, Kodama, & Igarashi, 1997). Recent studies have shown that Dnr also controls the expression of the H2-type VI secretion system (H2-T6SS) (Dang, Wang, Wen, & Liang, 2022), which contributes to metal ion acquisition, interbacterial competition, and biofilm fitness in low-oxygen environments (Han et al., 2019; Wang et al., 2021).

Although Dnr activation is NO-dependent, the optimal intracellular concentration of NO for Dnr function is poorly defined. Excessive NO may impair Dnr activity or other NO-sensitive regulators, potentially through disruption of iron-sulfur clusters or heme groups (Crack, Green, Hutchings, Thomson, & Le Brun, 2012; Crack, Stapleton, Green, Thomson, & Le Brun, 2013; Fontenot, Cheng, & Ding, 2022). Based on this rationale and the observation of disrupted expression of H2-T6SS in *norCBD* mutant, we investigated, in this study, whether *norCBD* disruption might impair Dnr-mediated gene expression by allowing toxic NO accumulation, thereby altering regulatory outputs, including the activation of H2-T6SS.

While the role of NorCBD in NO detoxification has been well documented (Zhou et al., 2019), its potential regulatory influence via modulation of intracellular NO levels and the resulting phenotypical changes have not been explored. We report the data suggesting that NorCBD not only facilitates anaerobic respiration but also contributes to the proper regulation of Dnr-dependent genes by maintaining intracellular NO balance that supports optimal Dnr activity.

## Materials and Methods

### Bacterial plasmids and strains

Bacterial strains and plasmids were used in all the experiments shown in **Table 1**. *E. coli* and *P. aeruginosa* were grown on Luria-Bertani (LB) agar and broth at 37°C. Different concentrations of antibiotics were applied. For *P. aeruginosa*, the concentration was carbenicillin (250 µg/ml), trimethoprim (300 µg/ml), and tetracycline (70 µg/ml). The antibiotic concentration used for *E. coli* was kanamycin (50 µg/ml), ampicillin (100 µg/ml), and tetracycline (12.0 µg/ml). Tetracycline (300 µg/ml) was used for constructing *P. aeruginosa* mutant PAO1(*ΔnorCBD*), which was plated on a Pseudomonas isolation agar (PIA) plate.

**Table 1.** List of bacterial strains and plasmids used.

| Bacterial strains and plasmids | Characteristics | Source/ Reference |
| --- | --- | --- |
| *P. aeruginosa* strains |  |  |
| PAO1 | Wild-type *P. aeruginosa* | Duan Lab |
| PAO1(Δ*norCBD*) | *norCBD* knockout mutant of PAO1 | This study |
| PAO1(Δ*clpV3*) | *clpV3* knockout mutant of PAO1 | (Li, Chen, Zhang, Bhagirath, & Duan, 2020) |
| *E. coli* strains |  |  |
| DH5α | F$^-$ Φ80lacZΔM15 $\Delta$(*lacZYA-argF*) U169<br>*recA1 endA1 hsdR17* (rK$^-$, mK$^+$) *phoA*<br>*supE44* $\lambda^-$ *thi-1 gyrA96 relA1* | Invitrogen |
| SM10-λ pir | Mobilizing strain, RP4 integrated into the chromosome; Km$^r$ | Duan Lab |
| Plasmids |  |  |
| pEX18Tc | *oriT*$^+$ *sacB*$^+$ gene replacement vector with multiple-cloning site from pUC18; Tc$^r$ | (Hoang et al., 1998) |
| pBT20 | Mini-TnM delivery vector; Gm | (Kulasekara et al., 2005) |
| pAK1900 | *E. coli*-*P. aeruginosa* shuttle cloning vector, Amp$^r$ | (Sharp, Jansons, Gertman, & Kropinski, 1996) |
| pMS402 | Expression reporter plasmid carrying the promoterless *luxCDABE*; Km$^r$ Tmp$^r$ | (Duan et al., 2003) |
| CTX-6.1 | Integration plasmid origins of plasmid mini-CTX- *lux*; Tc$^r$ | (Kong et al., 2013) |
| pRK2013 | Broad-host-range helper vector; Tra<sup>+</sup>, Km<sup>r</sup> | (Ditta, Stanfield, Corbin, & Helinski, 1980) |
| pEX18Tc-*norC* up | pEX18Tc carrying the upstream fragment of *norC* | This study |
| pEX18Tc-*norC*up+dw | pEX18Tc carrying the upstream and downstream fragment of *norC* | This study |
| pEX18Tc-*norD* up | pEX18Tc carrying the upstream fragment of *norD* | This study |
| pEX18Tc-*norD*up+dw | pEX18Tc carrying the upstream and downstream fragments of *norD* | This study |
| pAK-*norCBD* | pAK1900 with a 2874 bp fragment of *norCBD* between *KpnI* and *HindIII*; Amp<sup>r</sup>, Cb<sup>r</sup> | This study |
| CTX-*H2-T6SS* | Integration plasmid, CTX6.1 with a fragment of pKD-*hsiA2* containing *H2* promoter (pMS402 containing *hsiA2* promoter region) region and *luxCDABE* gene; Kn<sup>r</sup>, Tmp<sup>r</sup>, Tc<sup>r</sup> | This study |
| CTX-*H2-T6SS* in PAO1(Δ*norCBD*) | Integration plasmid, CTX6.1 with a fragment of pKD- *H2-T6SS*- PAO1(Δ*norCBD*) containing *H2* promoter region and *luxCDABE* gene; Kn<sup>r</sup>, Tmp<sup>r</sup>, Tc<sup>r</sup> | This study |
| CTX-*H2-T6SS* in PAO1(Δ*norCBD*)-*norCBD* in PAK1900 | Integration plasmid, CTX6.1 with a fragment of pKD- *H2-T6SS*- PAO1(Δ*norCBD*)-*norCBD* in PAK1900 containing *H2* promoter region and *luxCDABE* gene; Kn<sup>r</sup>, Tmp<sup>r</sup>, Tc<sup>r</sup> | This study |

### Construction of H2-T6SS transposon mutagenesis library

For making a transposon mutagenesis library, the CTX-H2-PAO1 strain was subjected to mutagenesis by using a vector known as pBT20, with some modifications in the methods as described (Kulasekara et al., 2005; Liang, Li, Dong, Surette, & Duan, 2008). In detail, PAO1 was used as a recipient strain, whereas *E. coli* SM10 containing pBT20 was used as a donor. First, the overnight culture of the donor and recipient was scraped from the culture plate, and the OD_600_ was adjusted to 40 and 20 for the donor and recipient, respectively. Donors and recipients were taken in an equal ratio (20μl). After mixing well, the culture was spotted on the LB agar plate and incubated at 37°C for 3 h. Then, mixed cultures were taken and diluted. After that, a PIA medium with gentamycin (Gm) (150 μg/ml) was used for spreading purposes. Around thirty thousand colonies were picked up and screened on 96-well plates, and a selective medium was used to make a transposon mutant library. Furthermore, colonies were incubated overnight in an LB medium containing selective antibiotics for the screening of genes that were involved in the expression of H2-T6SS. The next morning, the mutagenesis colonies were inoculated into 96-well plates, where the medium was LB (containing Gm at 50 μg/ml). For experimental purposes, both OD_600_ and luminescence were measured for 24 h at 37°C. Colonies with altered expression (two or more) were selected for further screening. To avoid false positive results, a total of six additional repeats were done. Colonies with altered H2-T6SS expressions were confirmed and further characterized.

To confirm the transposon insertion site, we conducted an arbitrary primed polymerase chain reaction (PCR), and DNA sequencing was done for the PCR products as described, with some modifications (Liang et al., 2008). In brief, two primers, P7-1, which read out from one end of the transposon, and the arb1 primer, arbitrary (**Table 2**), along with the chromosomal DNA of the transposon mutant, were used for the PCR template for the first round. In the second round, another two primers, P7-2 and arb2 (**Table 2**), were used for the PCR. After the PCR products were purified by the Gel/PCR Extraction Kit (Geneaid). PCR products were further sequenced at the Sequencing Core Facility of the Research Institute in Oncology and Hematology (RIOH), Manitoba. Finally, a BLAST query was done to confirm the transposon insertion site.

**Table 2.**
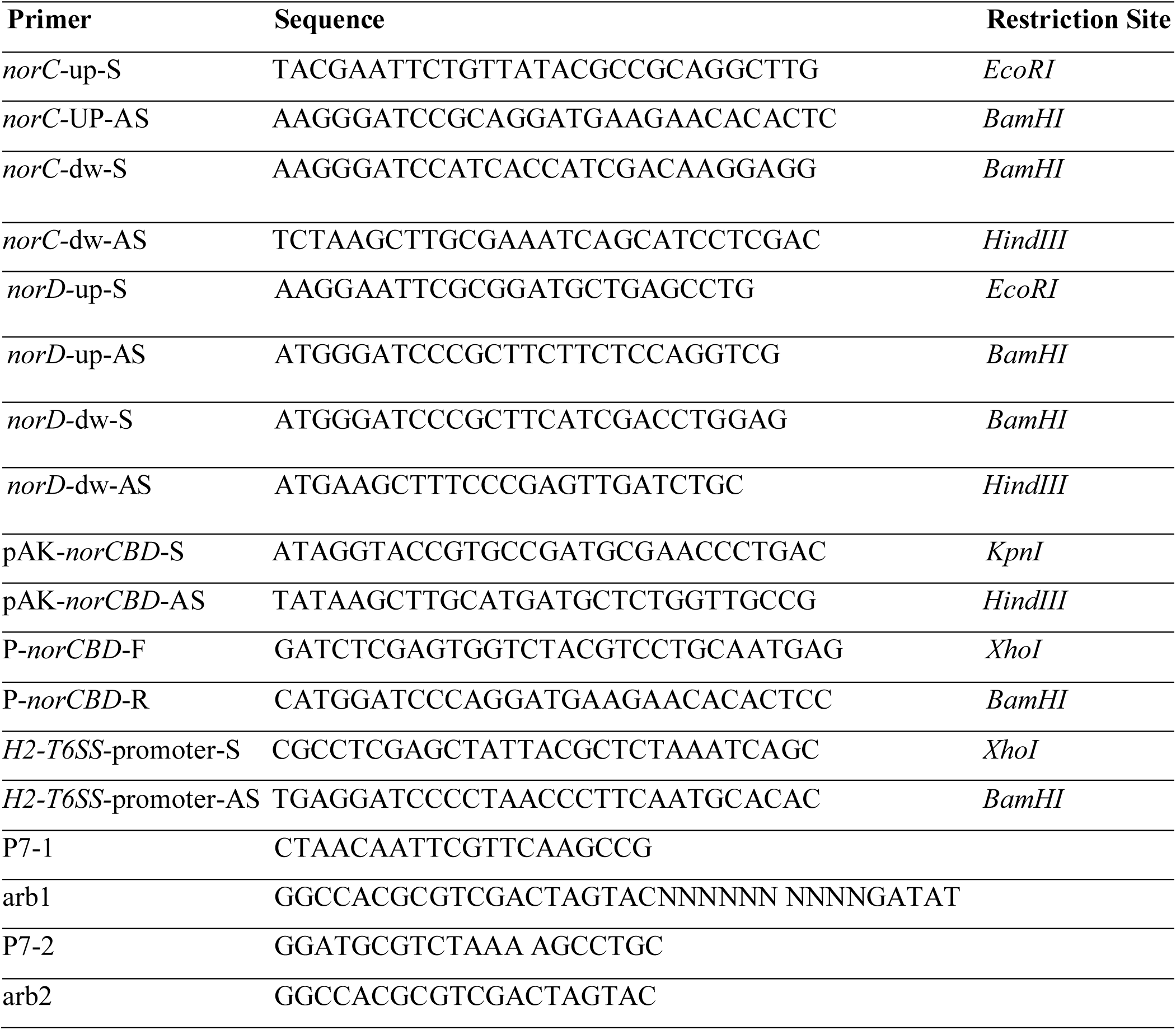
List of primers used.

### Generation of a *norCBD* deletion mutant PAO1 (*ΔnorCBD)* and a complementation strain

A deletion mutant of *norCBD* was generated in *P. aeruginosa* PAO1 using plasmid pEX18Tc (sucrose counter-selection system (Hmelo et al., 2015; Hoang, Karkhoff-Schweizer, Kutchma, & Schweizer, 1998). In brief, both up and downstream fragments of the gene of interest (*norC*) were amplified by polymerase chain reaction using the designed primers shown in **Table 2**. Then, amplified fragments of up and downstream were cloned into a pEX18Tc vector that contains a counter-selectable *sacB* gene.

After the double restriction enzyme digestion, both upstream and downstream products were ligated. PAO1(*ΔnorC*) was obtained by tri-parental mating. Triparental mating was performed using the *E. coli* (helper strain) that contains pRK 2013, the donor strain *E. coli* had pEX18Tc-*norC* (up and down), and the recipient strain used for this purpose was PAO1. Overnight cultures of the donor, helper, and recipient strains were resuspended in a PBS buffer. The bacteria were spotted on LB agar by mixing at a ratio of 2:2:1. An overnight culture was taken, and the developed pellets were resuspended in LB. The diluted bacteria suspension was spread on a PIA plate (containing tetracycline, 300 µg/ml) for the selection of merodiploid. For selecting a double crossover, the colony was streaked in LB agar containing 15% sucrose and no NaCl after the first crossover. Deletion mutants were verified by polymerase chain reaction using the confirmation primers **(Table 2).** Finally, PAO1(*ΔnorCBD*) was constructed using the same method as PAO1(Δ*norD*) as a background and was confirmed by the PCR.

Shuttle vector pAK1900 was used for the complementation of *norCBD* knockout mutants. First, the gene of interest (*norCBD*) was amplified using designed primers (pAK-*norCBD*-F and pAK-*norCBD*-R) (**Table 2**). Then, the amplified product was cloned into a shuttle vector, pAK1900. Finally, pAK-*norCBD* plasmid was transformed into PAO1(*ΔnorCBD*) by electroporation.

### Construction of a promoter-reporter-based gene expression detecting system

The plasmid pMS402 is used for constructing a promoter-*luxCDABE* reporter because this plasmid contains a promoterless *luxCDABE* reporter gene cluster (Duan, Dammel, Stein, Rabin, & Surette, 2003). First, the promoter region of *norCBD* will be generated by PCR using a primer (P-*norCBD*-forward and P-*norCBD*-reverse) designed for *norCBD* gene (**Table 2**) and inserted into BamHI-XhoI site of vector pMS402 (upstream of promoterless *luxCDABE*) to generate plasmid pKD-*norCBD*. Second, pKD-*norCBD* was digested with PacI, and the larger band was isolated by Gel DNA Fragment Extraction Kit and cloned into CTX6.1, which is also digested with PacI to generate plasmid CTX-*norCBD.* Then, CTX-*norCBD* was transformed into *E. coli* SM10-λ *pir.* Finally, the *P. aeruginosa* reporter integration strain CTX-*norCBD* in PAO1 was obtained by bi-parental mating (Hoang et al., 1998). In brief, the process was performed using the *E. coli* SM10-λ *pir* that contains CTX-*norCBD* and PAO1. Overnight cultures were resuspended in PBS buffer. The bacteria were spotted on LB agar by mixing at a ratio of 1:1. The Overnight culture was taken, and the developed pellets were resuspended in Super Optimal Medium with Catabolic Repressor (SOC) medium. The diluted bacterial suspension was spread on a PIA plate (containing tetracycline) for selection. CTX-*H2-T6SS* in PAO1 and CTX-*H2-T6SS* in PAO1(*ΔnorCBD)* were constructed using the method mentioned above.

### Measurement of promoter activities

Gene expression was measured by the lux-based reporter assays using a synergy plate reader (Biotek, USA) using the method as described (Duan et al., 2003). In detail, strains containing the reporter were grown overnight in liquid media (LB broth and M9 broth with supplements) and subcultured the next morning, and then diluted to an OD_600_ nm of 0.2. The diluted cultures were used as inoculants. After another 2 h of incubation, 5 μl of culture was inoculated into a 96-well flat-bottom plate containing 95 μl of fresh medium. A layer of 50 μl of filter-sterilized mineral oil (Sigma Aldrich) was applied on the surface to inhibit evaporation throughout the assay. Luminescence (counts per second, cps) was measured every 30 min for 24 h, and bacterial growth was monitored by measuring OD_600_ nm at the same time. The gene expression level was expressed by cps/OD_600_.

In the test of the effect of NO donor and NO scavenger on H2-T6SS expression, NO donor sodium nitroprusside (SNP) was added at different concentrations (10 µM, 25 µM, and 50 µM) in the media, and NO scavenger carboxy-phenyl-tetramethylimidazolineoxyl (CPTIO) (Cayman Chemical) was added at the concentrations of 2 mM, 5 mM, and 10 mM.

### Measurement of intracellular nitric oxide levels

The NO detection reagent diaminofluorescein-2 diacetate (DAF-2 DA) (Sigma-Aldrich, USA) was used to quantify the cellular NO levels of *P. aeruginosa* bacterial cells using a previously established method (Su et al., 2014). In detail, to reach the exponential growth phase (OD_600_ = 0.2), the bacterial strain was subcultured for 3.5-4 h after being incubated overnight at 37°C. Anaerobic bacterial growth was accomplished using the BD GasPak EZ Anaerobe Pouch System. The LB broth was supplemented with 50 mM KNO_3_ to promote anaerobic growth. We also measured the intracellular NO concentrations under normoxic conditions, where strains were cultured in a 96-well plate. In brief, when the OD_600_ nm reached around 0.2, a 96-well plate with a flat bottom was filled with 180 μl of fresh M9 minimal media supplement, along with 20 μl of bacterial cultures and 50 μl filtered (0.22μm) mineral oil, and allowed to grow inside the plate reader. Following anaerobic and normoxic growth (at 5 and 10 h), 1 ml of shaking culture was incubated with 10 µM DAF-2 DA for 1 h at 37°C. After incubation, the cell was washed with 1X PBS. Following washing, the reaction product DAF-2T’s fluorescence level was measured (an excitation wavelength of 495 nm and an emission wavelength of 515 nm), normalized to OD_600_ with the FlexStation 3 Multi-Mode Microplate Reader (Molecular Devices, San Jose, CA, USA).

### Pyocyanin production measurement

Overnight culture (18 h) was used to extract and quantify the pyocyanin production, and the methods were described previously (Essar, Eberly, Hadero, & Crawford, 1990). In brief, the overnight culture was centrifuged, and 5 ml of supernatant was mixed with 3 ml of chloroform. 1 ml of 0.2 N HCl was added to the chloroform layer and centrifuged at 4500 g for 10 min. After centrifugation, the upper layer (pink) was transferred to the cuvette, and absorbance was measured at 520 nm. The concentration of pyocyanin was expressed as micrograms produced per milliliter of culture supernatant, calculated by multiplying the extinction coefficient of 17.072 at 520 nm.

### Measurement of biofilm formation

Biofilm formation was measured by the protocol as described previously (O’Toole, 2011). Overnight cultured cells were inoculated into M63 medium (1:100 dilutions) containing glucose (0.2%), casamino acids (0.5%), and MgSO_4_ (1 mM), and 96-well microtiter plates were incubated at 37°C for 24 h. The planktonic cell was removed after washing with 1X PBS (three times). Crystal violet solutions (100 μl) were added, and staining was allowed for each well at room temperature (15 min). All the wells were rinsed again with distilled water. 125 μl acetic acid (30%) in water was dissolved in the remaining crystal violet on the plate. Finally, 100 μl solution was transferred into a new 96-well plate, and absorbance was measured at 550 nm.

### Detection of proteolytic activity

Proteolytic activity was measured using the method known as agar plate assay, as mentioned in the earlier studies, with some modifications (Gupta, Gobble, & Schuster, 2009). Two microliters of cells from the overnight culture were plated on LB agar with 2% skim milk and incubated at 37°C overnight. The presence of clear zones around the bacterial colonies suggests proteolytic activity, and the diameters of these zones were recorded.

### Motility assays

Bacterial motility activities were assessed according to the previously described protocol (Rashid & Kornberg, 2000). The medium for swarming motility contained glucose (5 g/l), agar (0.5%), and nutrient broth (8 g/l). The medium for swimming motility contained agar (0.3%), NaCl (5 g/l), and tryptone (10 g/l). Bacteria were spotted with 2 μl of the aliquot of the overnight LB culture for swarming and swimming motility assays. For swimming motility, the incubation was 15 h at 37°C. For swarming motility, the incubation was at room temperature for 8 h, followed by incubation at 37°C for 15 h. After inoculation, photographs were taken with a Vilber Lourmat Imager (Fusion FX7).

### Intracellular cAMP levels quantification

The intracellular cAMP levels were quantified using the cAMP Select ELISA kit (Cayman Chemical, California, US). The cells from an overnight culture (OD_600_ was adjusted to 8) were collected by centrifugation, and 5 ml of culture was mixed with 500 μl of 0.1N HCl. The cells were allowed to undergo 30 s sonication (120 V). The bacterial cells were then centrifuged at 1000 g for 10 min. After centrifugation, the supernatant was collected. The bacterial sample and standard were prepared according to the manufacturer’s protocol and loaded into a 96-well plate supplied by the manufacturer. The 96-well plate was then covered with a plastic film and allowed for orbital shaking under dark conditions (for 90 min). Using a plate reader, the absorbance was read at around 405 and 420 nm. The optical density of the bacterial sample concentration was determined from the standard curve plot using the equation as per the manufacturer’s guidelines.

### Statistical analysis

All experimental results are presented as the mean ± Standard error of the mean (SEM) from three independent experiments. Statistical significance tests and analysis were performed by using GraphPad PRISM version 10 (GraphPad Software, San Diego, CA, USA). Two-way analysis of variance (ANOVA) was applied to compare more than three groups with two independent variables. One-way ANOVA was used to compare more than three groups with one independent variable. Wherever applicable, significance levels are indicated as follows: ns (not significant), *p* > 0.05; * *p* < 0.05, ** *p* < 0.01, *** *p* < 0.001, and **** *p* < 0.0001.

## Results

### Screening of genes affecting H2-T6SS by transposon mutagenesis

*P. aeruginosa* has three homologous yet functionally distinctive T6SS systems, H1-T6SS, H2-T6SS, and H3-T6SS, with H1-T6SS being the most studied. In the search for H2-T6SS regulators, a transposon mutant library was constructed to identify the regulatory pathway of T6SS. A transposon mutagenesis was performed in PAO1 harboring the CTX-H2-T6SS*-lux* reporter integrated on the chromosome by utilizing the mariner transposon vector pBT20 as previously described (Kulasekara et al., 2005; Liang et al., 2008). Approximately 30,000 colonies grown on selective media containing gentamicin were screened for altered H2-expression. Colonies with changed light production under an imager were selected for further studies. Arbitrary PCR and sequencing were performed to identify the transposon insertion sites in these mutants, as previously reported (Liang et al., 2008). From this transposon mutant library, eight genes were identified whose transposon insertion affected the expression of H2-T6SS (**Table 3**). Among them, PA0523 (*norC*) and PA0525 (*norD*) are located in the same operon. *norCBD* operon encodes nitric oxide reductase (NorCBD), which reduces nitric oxide (NO) to nitrous oxide (N₂O), a key step for reducing the toxic level of intracellular NO and anaerobic respiration (Arai, 2011; Hino et al., 2010).

**Table 3:** Potential regulators of *H2*-T6SS (Transposon mutants)

| Gene name or number | Insertion site | Protein description | Max Fold<sup>a</sup> |
| --- | --- | --- | --- |
| *PA0523(norC)* | 582172 | Nitric oxide reductase subunit C | -2.8 |
| *PA0525 (norD)* | 585588 | Probable denitrification protein | -2.8 |
| *PA0337(ptsP)* | 380328 | Phosphoenolpyruvate-protein<br>phosphotransferase ptsp | +2.5 |
| *PA0961-0962* | 1046969 | Probable cold-shock protein | +3 |
| *PA1689* | 1840252 | Conserved hypothetical protein | +3 |
| *PA2929* | 3285258 | Hypothetical protein | -2 |
| *PA3798* | 4257397 | Probable aminotransferase | -2.5 |
| *PA4467* | 4997059 | Hypothetical protein | -2 |

<sup>a</sup> Max fold, the maximal ratio of expression between the mutant and the wild-type

### *norCBD* disruption affects the expression of H2-T6SS

To verify the effect of *norCBD* disruption on the H2-T6SS expression, we constructed a *norCBD* deletion mutant PAO1(Δ*norCBD*) and compared the promoter activities of H2-T6SS in PAO1 and in the mutant. Same as the PAO1 harboring the CTX-H2-T6SS*-lux* reporter, the CTX-H2-T6SS*-lux* reporter was also chromosomally integrated in PAO1(Δ*norCBD*). A complementation strain of the mutant carrying the entire *norCBD* on a plasmid was also constructed. The transcriptional activities of H2-T6SS were measured and compared in these strains by measuring luminescence. Our results show that deletion of *norCBD*, significantly reduced the CTX-*H2*-T6SS transcriptional activity (**Figure 1**), confirming the involvement of *norCBD* in the regulation of H2-T6SS. The maximal promoter activity of H2-T6SS at 8 hrs was around 8-fold lower in PAO1(Δ*norCBD*) compared with that in the wild-type PAO1. The complementation of *norCBD* restored the expression of H2-T6SS to the wild-type PAO1 level.

**Figure 1.**
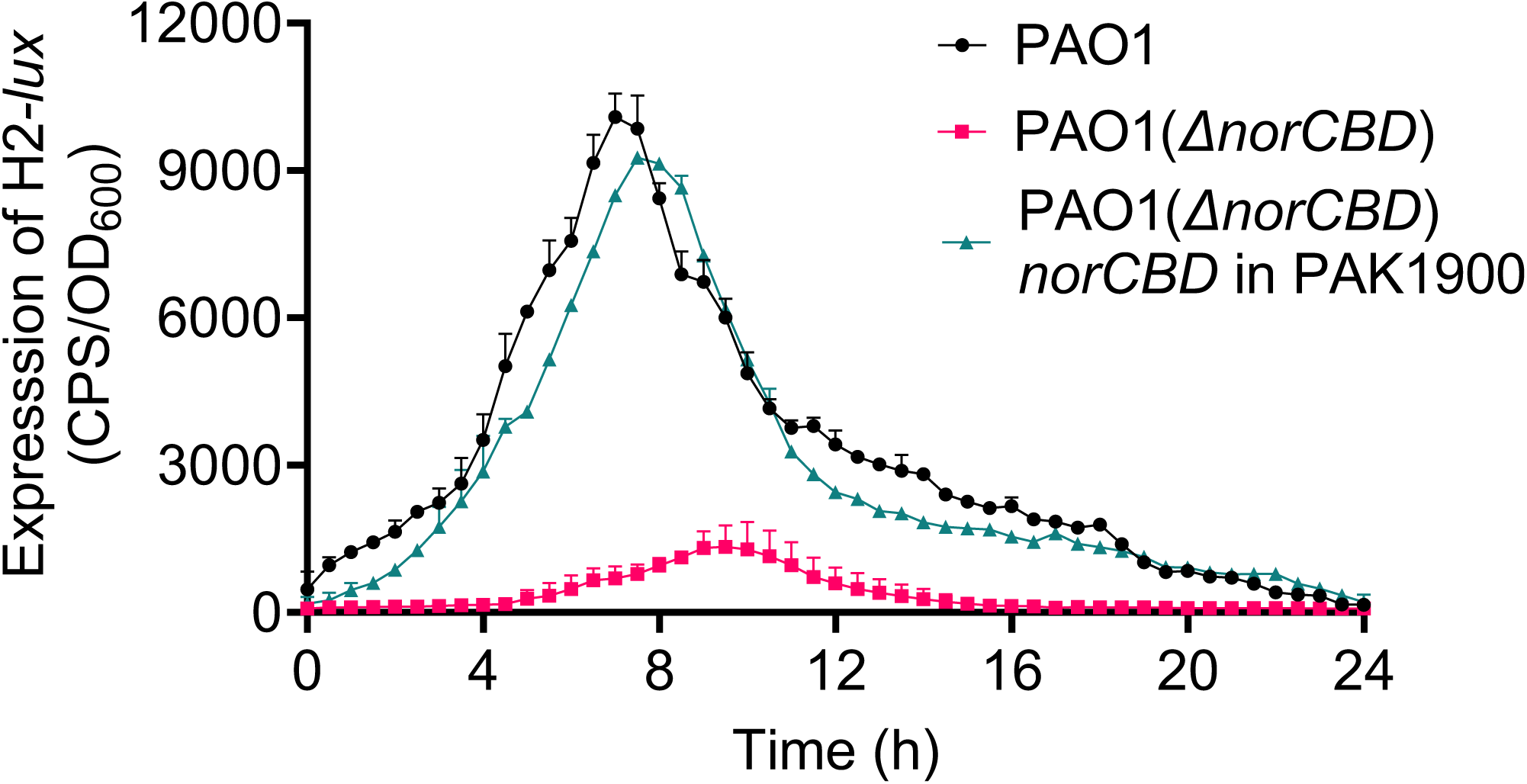
Deletion of *norCBD* resulted in decreased expression of H2-T6SS. The CTX-H2-T6SS reporter fusion integrated on the chromosome was used to measure the promoter activity of the H2-T6SS gene in PAO1(*ΔnorCBD*). The result represents the average of the triplicate experiments, and the error bars indicate standard deviations. The result showed the decreased expression of the H2 reporter under PAO1(*ΔnorCBD*). When comparing PAO1(*ΔnorCBD*) to the wild-type PAO1, the promoter activity of H2-T6SS was significantly lower in the deletion mutant.

### Effect of NO donor and NO scavenger on the H2-T6SS expression

The *norCBD* operon encodes nitric oxide reductase (NorCBD), which reduces nitric oxide (NO) to nitrous oxide (N₂O). It is known that H2-T6SS is activated by Dnr in response to NO (Dang et al., 2022). It is expected that disruption of norCBD could increase the level of NO, which in turn would increase H2-T6SS expression. However, our results showed the opposite effect, namely, that potentially increasing NO in the mutant actually repressed H2-T6SS expression.

To test the possibility that the altered NO levels caused by the disruption of *norCBD* somehow contributed to the observed decrease of H2-T6SS expression in the *norCBD* mutant, we examined the expression of H2-T6SS in PAO1, PAO1(*ΔnorCBD),* and the complementation strain in the presence and absence of NO donor, -SNP.

As shown in **Figure 2**, the *H2*-T6SS expression was significantly changed following the addition of the SNP. However, opposite changes were observed in the *norCBD* mutant compared to the wild type and the complementation strain. The expression of *H2*-T6SS was significantly increased in wild-type PAO1 compared to that in the absence of SNP (**Figure 2A**). The complementation strain of *norCBD* showed similar results to wild-type PAO1 (**Figure 2C**). In the *norCBD* deletion mutant, PAO1(*ΔnorCBD),* a significant reduction in H2-T6SS expression was observed in the presence of SNP (**Figure 2B**). This observation contrasts with the increased *H2*-T6SS expression observed in the wild type and complementation strains. These results appear to support our hypothesis that altered NO caused by *norCBD* disruption contributes to the changed H2-T6SS expression. In the wild type and the complementation strain, the increased H2-T6SS expression is probably a result of elevated Dnr activation in response to the increased NO level in the presence of SNP. In the norCBD mutant, however, further increase of NO on the already accumulated level potentially further impaired the Dnr-mediated H2-T6SS activation.

**Figure 2.**
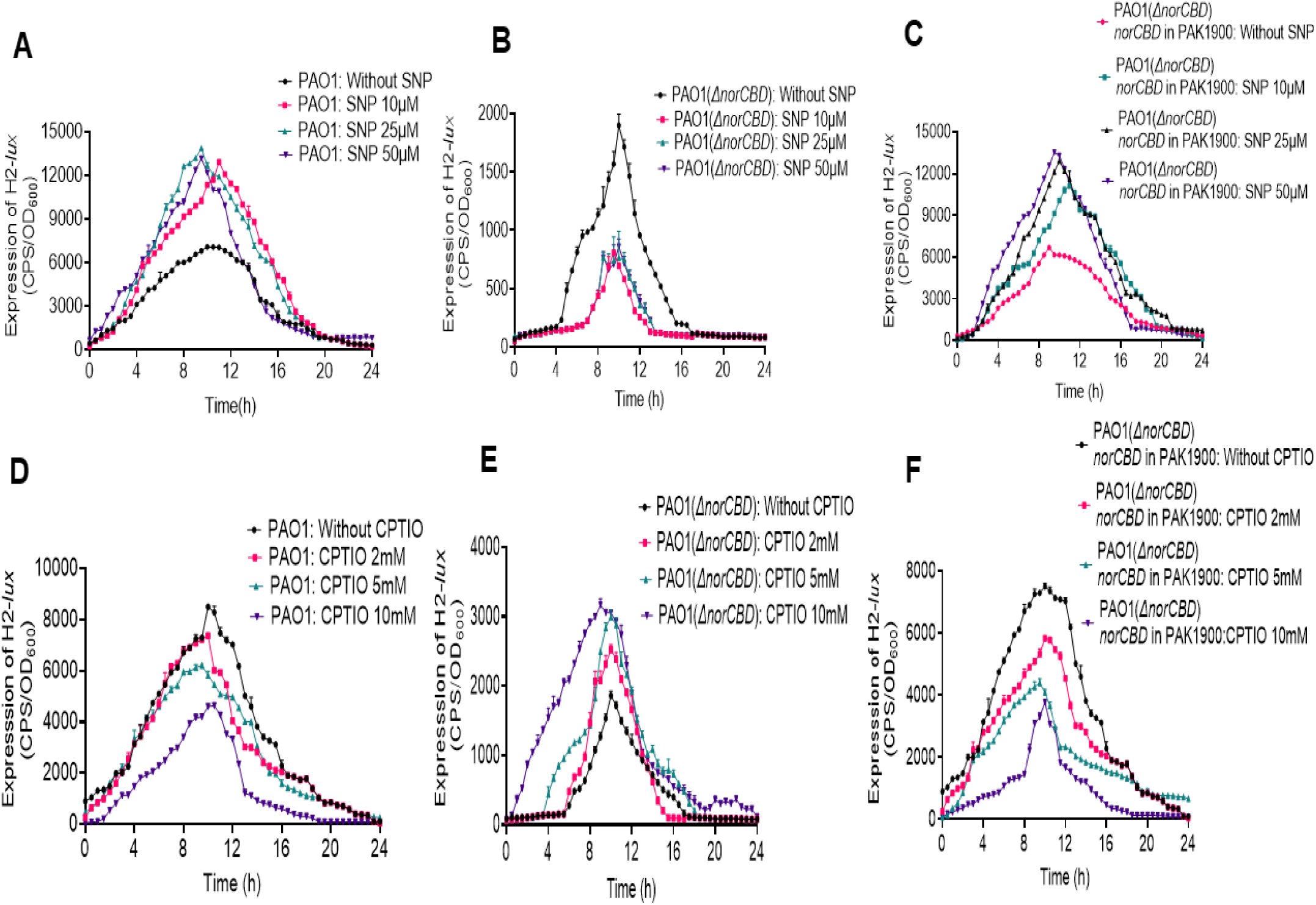
Effect of NO donor SNP and NO scavenger CPTIO on H2-T6SS expression: In this experiment, different concentrations of SNP (10 µM, 25 µM, and 50 µM) and CPTIO (2 mM, 5 mM, and 10 mM) were used. The expression of H2-T6SS was significantly affected by different concentrations of SNP and CPTIO. **(A)** Our findings from this 24 h experiment demonstrated that the highest expression of H2-T6SS in PAO1 was achieved after 10 h, with the highest expression of the SNP at a dose of 25 µM, indicating the use of NO. **(B)**When compared to the expression level without SNP, H2-T6SS expression was considerably lower for the PAO1(*ΔnorCBD*) strain. In comparison to the control, the lowest expression level was attained at a concentration of 50 µM SNP after 10 h. **(C)** Similar outcomes to those of the wild-type PAO1 were likewise displayed by the complementation strain of norCBD combined with CTX-H2-T6SS. Additionally, the experiment demonstrated that at a concentration of 25 µM SNP, the expression level peaked at 10 h. **(D)** Over time, the outcomes in wild-type PAO1 demonstrated a dose-dependent pattern. The expression of H2-T6SS gradually dropped when we raised the concentration of CPTIO to 5 mM and 10 mM, indicating that nitric oxide is being scavenged in the wild-type strains. When the H2-T6SS was examined, especially after 10 h, it was discovered that, in comparison to the control, the concentration of 10 mM revealed a low level of wild-type expression. **(E)** In the PAO1(*ΔnorCBD*), when we increased the CPTIO concentration, the expression level increased considerably. After 10 h, the maximum expression was observed at CPTIO concentrations of 5 mM and 10 mM. **(F)** The complementation strain’s H2-T6SS expression level coincides with that of the wild-type. Error bars indicate standard deviations.

To further pursue this line of inquiry, the effect of NO scavenger -CPTIO on the H2-T6SS expression was also determined. The transcriptional activity of CTX-H2-T6SS was similarly assessed in the wild-type PAO1, PAO1(*ΔnorCBD)*, and complementation strains in the presence and absence of NO scavenger CPTIO. CPTIO reacts rapidly and stoichiometrically with NO to form nitrogen dioxide (NO₂), effectively removing NO from the cell (Goldstein, Russo, & Samuni, 2003; Lichtenberg et al., 2021). As shown in **Figure 2**, with CPTIO, the expression level of H2-T6SS was significantly reduced in wild-type PAO1 (**Figure 2D**) and in the complementation strain of *norCBD* (**Figure 2F**) in a dose-dependent manner. As anticipated, H2-T6SS expression was elevated in the PAO1(*ΔnorCBD* upon treatment with CPTIO (**Figure 2E**), indicating that NO scavenging alleviates the inhibition of Dnr activation caused by NO accumulation in the mutant.

### Accumulation of NO in the *norCBD* deletion mutant

Nitric oxide (NO) is a major signaling molecule in *P. aeruginosa,* but also toxic to the cells at high levels (Bogdan, Rollinghoff, & Diefenbach, 2000; Nisbett & Boon, 2016). To confirm the altered NO levels caused by the disruption of *norCBD*, we measured the level of intracellular NO using diaminofluorescein-2 diacetate (DAF-2 DA), which is a cell-permeable fluorescent NO indicator converted to DAF-2 by the intracellular esterases. Upon reacting with nitric oxide, DAF-2 produces DAF-2 TA, a derivative that can be measured by fluorescence. The fluorescent intensity of DAF-2 TA gives a real-time measurement of NO. As expected, the *norCBD* deletion mutant showed significant accumulation of NO compared to the wild-type PAO1(**Figure 3A**) under anaerobic conditions. At the early exponential phase (5 h), NO in the *norCBD* mutant was 4.37-fold higher than that in the wild-type PAO1. The complementation strains showed the accumulation of nitric oxide, which is close to wild-type PAO1. The difference in NO levels between the mutant and the wild type persisted, although the NO levels in both strains were lower than those in the exponential stage.

**Figure 3.**
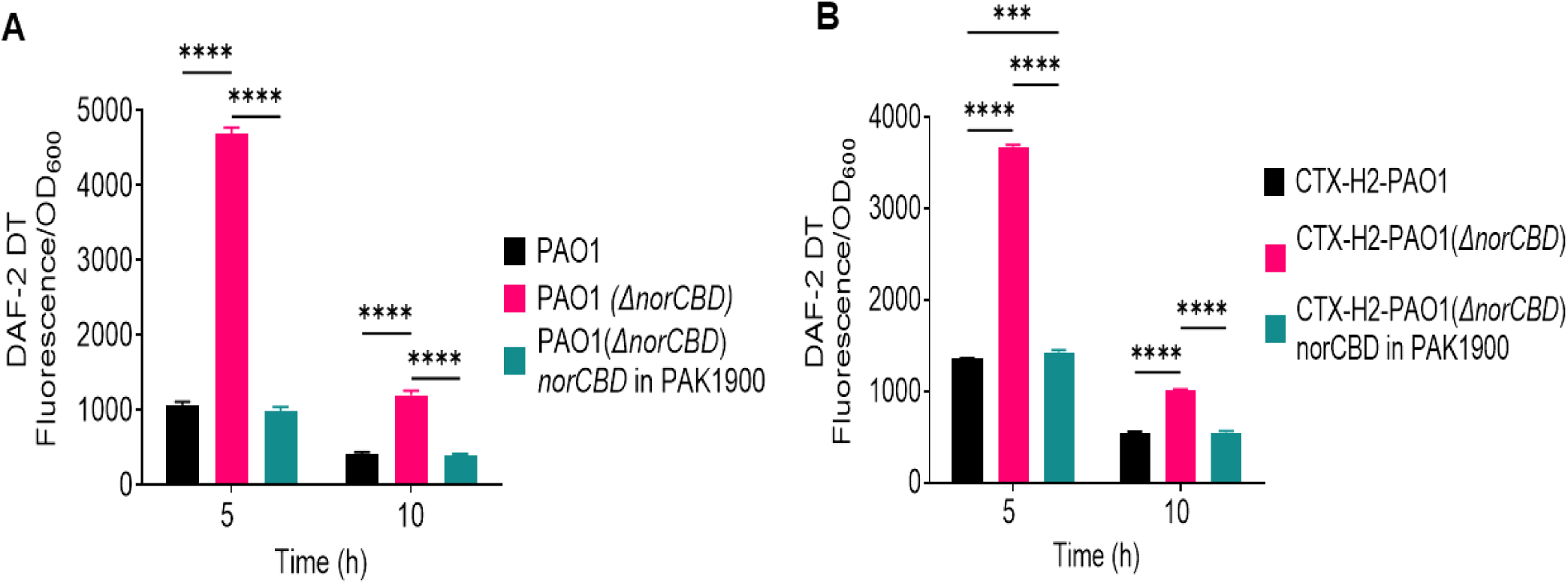
Intracellular nitric oxide was accumulated in the *norCBD* deletion mutant. The level of nitric oxide was measured in wild-type PAO1, PAO1(*ΔnorCBD*) mutant, and its complemented strain using the NO detection reagent diaminofluorescein-2 diacetate (DAF-2 DA). Within the cell, esterases transform the cell-permeable molecule DAF-2 DA into DAF-2. A fluorescent triazole derivative (DAF-2T) is formed when DAF-2 and NO react with each other. The amount of cellular NO present is directly proportional to the increase in DAF-2TA fluorescence intensity, as measured at a particular wavelength. At each time point, measured fluorescence intensity was normalized to OD_600_. The initial accumulation of intracellular nitric oxide (real-time) was observed at two points, 5 h and 10 h. **(A)** Under anaerobic conditions, at 5 h and 10 h, the intracellular accumulation of nitric oxide in PAO1(*ΔnorCBD*) was respectively 4.37 and 3.06-fold higher when compared with the wild type PAO1. **(B)** Under normoxic conditions, nitric oxide still accumulates in higher amounts (2.71 and 1.87-fold in 5 h and 10 h, respectively) in CTX-H2-PAO1(*ΔnorCBD*) compared with the wild-type CTX-H2-PAO1. Overall, the accumulation of nitric oxide in PAO1(*ΔnorCBD*) was relatively higher under anaerobic conditions. Data were analyzed using two-way ANOVA and Tukey’s multiple comparisons test. Error bars indicate standard deviations. *** *p* < 0.001 and **** *p* < 0.0001.

As most of our H2-T6SS expression was measured in normoxic conditions, we also measured the intracellular NO concentrations in such conditions where the strains were cultured in a 96-well plate. Similarly, the results obtained (**Figure 3B**) showed that at 5 h and 10h, the intracellular NO level in the *norCBD* mutant was significantly elevated (2.71-fold and 1.87-fold, respectively) compared to the wild type and the complementation strains. The results confirm that NorCBD disruption causes an unchecked increase of intracellular NO, which likely results in dysregulation of H2-T6SS.

### Bacterial growth affected by *norCBD* disruption and NO accumulation

It was noted during the NO measurement that PAO1(*ΔnorCBD*) exhibited reduced growth compared to wild-type PAO1 and the complementation strain (**Figure 4A**) under anaerobic conditions. We reasoned that the accumulation of NO and disruption of anaerobic respiration in the *norCBD* mutant affected the growth. To further investigate the impact of NO accumulation and denitrification disruption on the growth of the *P. aeruginosa*, we compared the growth of norCBD mutant with that of the wild type and complementation strain under normoxic conditions (**Figure 4B**), and we also examined the effects of both an NO donor (**Figure 4C**) and an NO scavenger (**Figure 4D**) to determine whether modulation of NO levels could influence growth outcomes. As shown in **Figure 4**, the growth of the *norCBD* mutant consistently was lower than that of the wild type or the complementation strains, suggesting the importance of intracellular NO balance and functional denitrification metabolic pathway in *P. aeruginosa*.

**Figure 4.**
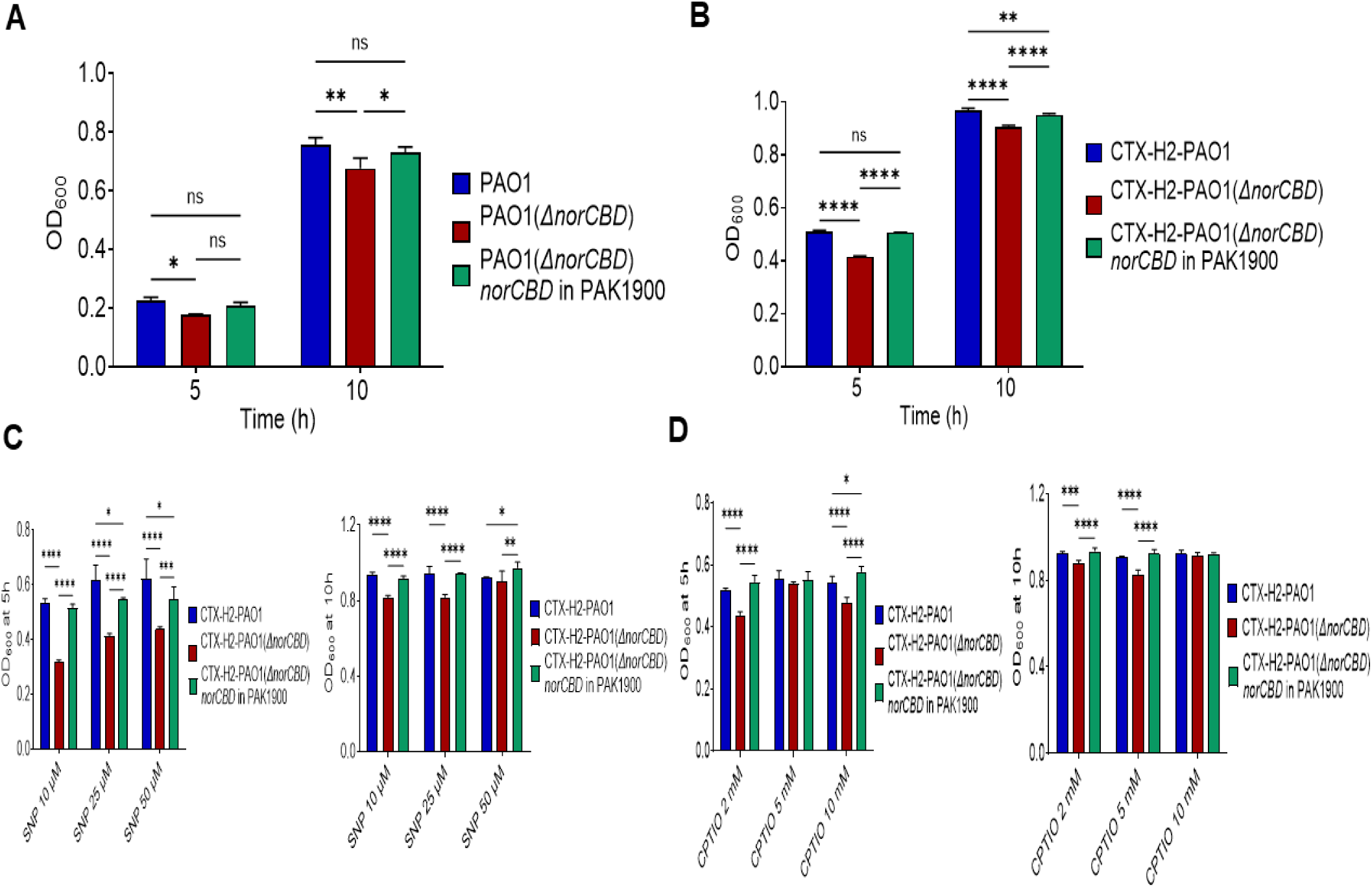
Characterization of *P. aeruginosa* growth (OD_600_) under different conditions. The growth of wild-type PAO1, PAO1*(ΔnorCBD),* and complementation strains, PAO1*(ΔnorCBD)-*norCBD in PAK1900, was observed at 5 h and 10 h under anaerobic conditions, normoxic conditions, and with the addition of NO donor, SNP, and NO scavenger, CPTIO. **(A)** Under anaerobic conditions, the growth is slower in wild-type PAO1, PAO1*(ΔnorCBD),* and complementation strains compared with aerobic conditions compared to the normoxic conditions. **(B)** Under normoxic conditions, the OD_600_ was relatively higher in all strains, in contrast to the anaerobic growth conditions. (**C, D**) At 5 h and 10 h, growth (OD_600_) was monitored with the addition of different concentrations of SNP (10 µM, 25 µM, and 50 µM) and CPTIO (2 mM, 5 mM, and 10 mM). Overall, in different growth conditions, PAO1*(ΔnorCBD)* showed lower growth compared to the wild-type PAO1 and the complementation strain. Data were analyzed using two-way ANOVA and Tukey’s multiple comparisons test. Error bars indicate standard deviations. *ns* (not significant) *p* > 0.05; * *p* < 0.05, ** *p* < 0.01, *** *p* < 0.001, and **** *p* < 0.0001.

### Altered virulence traits in the *norCBD* mutant

NO serves as a signaling molecule in *P. aeruginosa* (Michaelis et al., 2024). It reduces intracellular levels of cyclic di-GMP, a second messenger that promotes biofilm formation. and influences quorum-sensing systems (Barraud et al., 2009; Plate & Marletta, 2012). NO is endogenously produced in the denitrification pathway, linking it to metabolic and environmental sensing functions. To assess the broader effect of *norCBD* disruption, we tested several phenotypes in the *norCBD* mutant strain PAO1(Δ*norCBD*), together with PAO1 and complementation strains. The phenotypes tested include pyocyanin production, biofilm formation, proteolytic activity, and motility.

As shown in **Figure 5**, when *norCBD* was inactivated, the level of pyocyanin was significantly reduced. The complementation of *norCBD* on a plasmid was able to restore the pyocyanin production to the wild-type level (**Figure 5A**). The biofilm formation assay showed that the PAO1(Δ*norCBD*) had a lower ability to form biofilms compared to the wild-type PAO1 (**Figure 5B**). Similarly, the *norCBD* mutant also showed reduced proteolytic activity (**Figure 5C**) and bacterial motility, especially the swimming motility and swarming motility (**Figure 5D).** These findings indicate that while NO is an important signal to activate multiple virulence-related phenotypes such as pyocyanin production, biofilm formation, proteolytic activity, and motility, the accumulation of NO in the PAO1(Δ*norCBD*) probably overrides its signaling function and results in the disruption of protease and pyocyanin production, biofilm formation, and motility.

**Figure 5.**
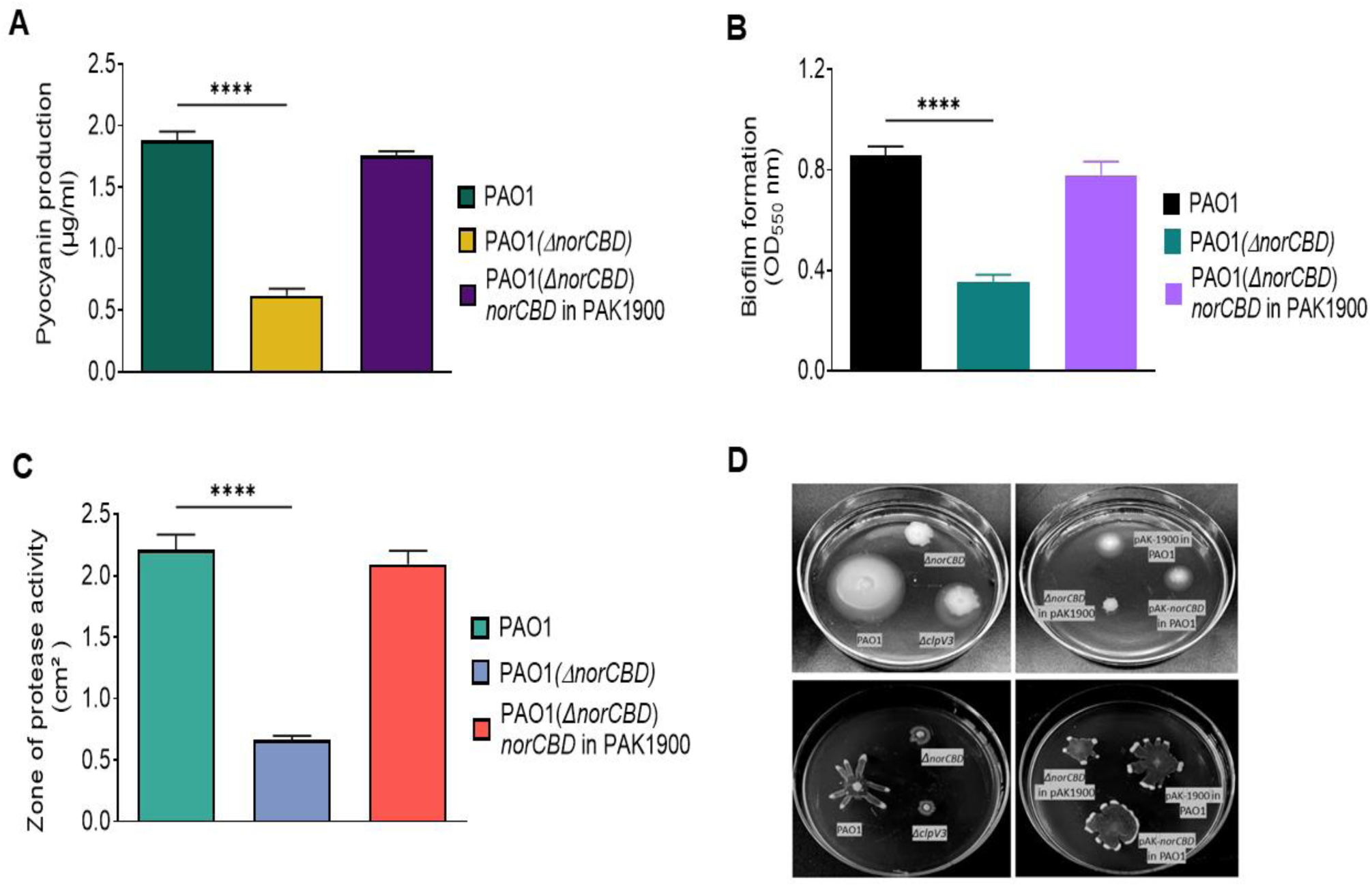
Multiple phenotypes of *P. aeruginosa* were affected by the deletion of the *norCBD* gene. **(A, B, C)** Pyocyanin production, biofilm formation, and proteolytic zone were significantly decreased in PAO1(*ΔnorCBD*) compared with PAO1 or the complementation strain. (**D**) Inactivation of *norCBD* in *P. aeruginosa* results in reduced swimming and swarming motility in PAO1(*ΔnorCBD*) compared with PAO1 or the complementation strain. Data was analyzed using one-way ANOVA. Error bars indicate standard deviations. **** *p* < 0.0001.

### Deletion of the *norCBD* gene reduced levels of the second messenger cAMP

It has been shown that the expression of H2-T6SS is positively regulated by NO-Dnr (Dang et al., 2022) but negatively modulated by cAMP-Vfr in *P. aeruginosa*. To investigate whether the observed phenotypical changes in the *norCBD* knockout mutant could be a result of altered cAMP levels, we compared the level of cAMP in the *norCBD* mutant with that of the wild type and the complementation strains. Using the commercially available Cyclic AMP Select ELISA Kit, we assessed the intracellular concentrations of cAMP in PAO1 and PAO1(Δ*norCBD*), and the results obtained indicate that the disruption of *norCBD* actually reduced the cAMP level (**Figure 6)** when compared with the wild-type PAO1 and the complementation of *norCBD*. Considering the previous report that the expression of T6SS inversely correlates with cAMP levels (Zhang et al., 2022), these results couldn’t explain the decreased H2-T6SS expression observed in the *norCBD* mutant. It is possible that the effect of the NO imbalance on H2-T6SS overridden the cAMP effect in the mutant.

**Figure 6.**
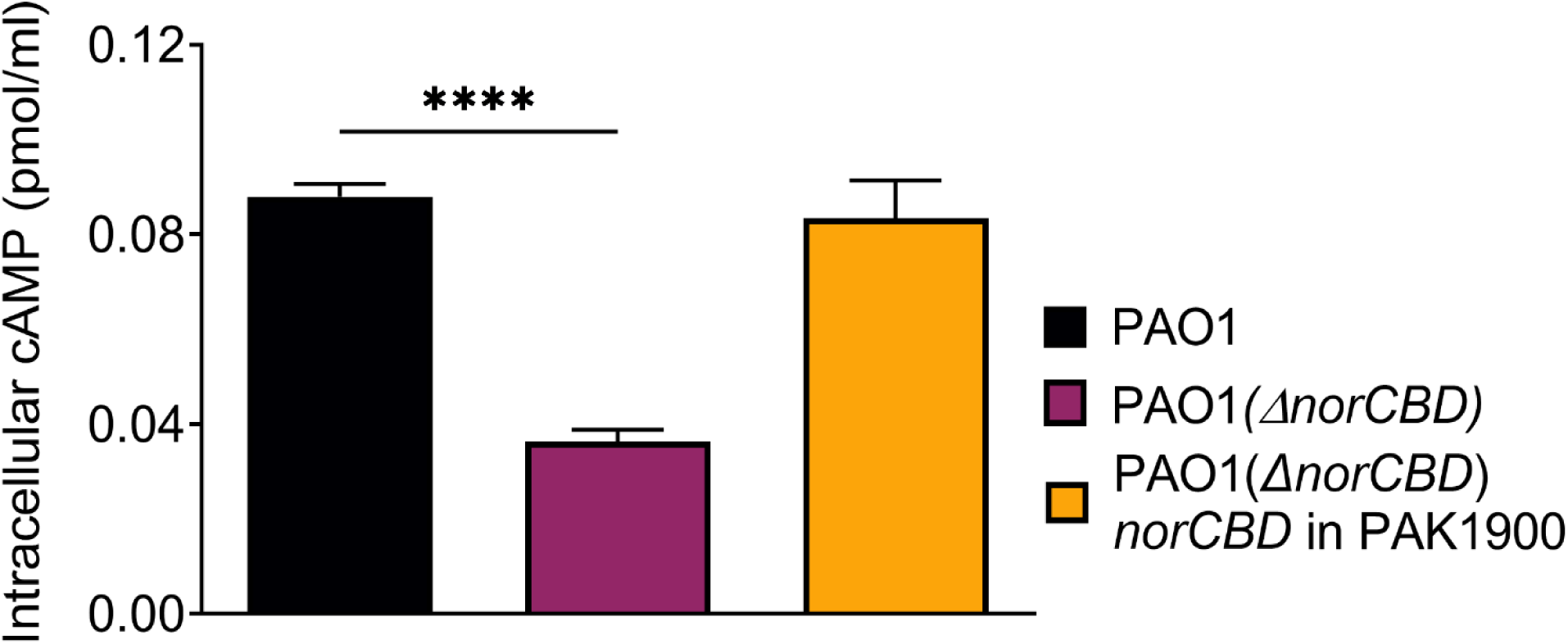
PAO1(*ΔnorCBD*) had reduced levels of second messengers, cAMP. The intracellular concentration of secondary messenger, cAMP, was significantly reduced when the *norCBD* gene was deleted. cAMP concentration was quantified in wild-type PAO1, PAO1(*ΔnorCBD*) mutant, and its complemented strain using the Cyclic AMP Select ELISA Kit. Data was analyzed using one-way ANOVA. Error bars indicate standard deviations. **** *p* < 0.0001.

## Discussion

This study identifies the *norCBD* operon as a previously unrecognized component that positively influences H2-T6SS transcription in *P. aeruginosa*. Deletion of *norCBD* elevated intracellular nitric oxide (NO) yet paradoxically suppressed H2-T6SS promoter activity; expression was restored by genetic complementation and by NO scavenging with CPTIO and was enhanced by an NO donor (SNP) only in wild-type and complemented strains. The *norCBD* mutant showed impaired growth, most evident under denitrifying conditions and broad defects in pyocyanin production, proteolysis, biofilm formation, and motility. Together, these observations support a model in which NorCBD maintains an optimal intracellular NO range required for Dnr-dependent activation of H2-T6SS and associated virulence programs; excessive NO pushes cells outside this range, attenuating Dnr output and disrupting multiple traits.

The biphasic behavior of NO on H2-T6SS links two seemingly opposing effects: NO is the activating signal for the Anr–Dnr–H2-T6SS cascade (Luo et al., 2025; Wang et al., 2021), yet too much NO is inhibitory. In our data, moderate NO in the wild type enhances H2-T6SS, whereas further NO elevation in the *norCBD* mutant depresses expression. A possible explanation is that high NO nitrosylates or otherwise damages NO-sensitive cofactors such as Fe–S clusters or heme groups in Dnr and other regulators (Crack et al., 2013; Mettert & Kiley, 2014), reducing their transcriptional activity despite NO being the initial activating signal. The rescue of H2-T6SS by CPTIO in the mutant, and the enhancement by SNP when NorCBD is present to balance NO concentration, are consistent with this interpretation. The growth defect of the *norCBD* knockout mutant under anaerobic conditions is consistent with the enzyme’s canonical role as the terminal NO reductase of denitrification (Arai, 2011; Zumft, 1997); without NorCBD, NO accumulates, respiration is compromised, and the resulting stress spreads into regulatory circuits that shut down non-essential systems, including T6SS.

Prior works have established that Anr and Dnr activate H2-T6SS genes and that H2-linked effectors promote fitness and competition in low-oxygen niches (Luo et al., 2025; Wang et al., 2021). Our results corroborate this Dnr-dependent logic but add an upstream physiological layer: endogenous NO homeostasis, maintained by NorCBD, is necessary for this pathway. The relevant variable appears to be not NO presence per sc, but its intracellular range. This agrees with broader observations that redox and nitrosative states interface with multiple global regulators in *P. aeruginosa* (Barraud et al., 2009; Korner, Sofia, & Zumft, 2003), including those that control acute–chronic transitions and metal acquisition, and suggests that H2-T6SS integrates respiratory status more tightly than previously appreciated.

Our data also extend the role of NorCBD beyond classical anaerobic metabolism. A recent report by Stuut Balsam *et al*. (2025) demonstrated that Dnr contributes to fitness not only under strictly anoxic conditions but also under microoxic and even normoxic conditions in the presence of low micromolar nitrate, as is found in standard LB medium (Stuut Balsam, Conaway, Mould, Jean-Pierre, & Hogan, 2025). In that study, nitrate consumption and NO formation under microoxia, and in LasR⁻ strains even under normoxia, created a dependence on Dnr-regulated NorCBD for detoxification and survival. Our findings align with this conclusion by showing that NorCBD is required to buffer NO and maintain optimal Dnr activity under both normoxic and oxygen-limited conditions. This positions NorCBD not only as a denitrification enzyme but also as a regulator of NO signaling in environments with fluctuating oxygen and nitrate availability, such as the cystic fibrosis lung, where LasR⁻ strains are commonly encountered and nitrate is abundant in sputum (Kolpen et al., 2014). Thus, NorCBD’s influence extends into normoxic physiology, providing a mechanism that safeguards Dnr-dependent transcriptional programs against NO toxicity in diverse environmental contexts. Such a model is consistent with the fact that *norCBD* is under direct transcriptional control of Dnr in *P. aeruginosa*, further reinforcing its role as both a metabolic enzyme and a regulator of NO-dependent signaling.

The changes in virulence-related traits observed in our study further support network-level coupling between denitrification physiology and global regulatory pathways. NO is known to reprogram second-messenger signaling; in particular, mis-timed or excessive NO can blunt c-di-GMP–driven switches that underlie biofilm formation and motility (Barraud et al., 2009; Li, Heine, Entian, Sauer, & Frankenberg-Dinkel, 2013). We also observed a decrease in intracellular cAMP in the *norCBD* mutant. Although the cAMP–Vfr signaling axis coordinates diverse virulence-related phenotypes, including T6SS in *P. aeruginosa* (Fuchs et al., 2010; Zhang et al., 2022), the reduction in cAMP observed in the *norCBD* mutant does not explain the decreased expression of H2-T6SS genes. One possibility is that the dysregulation of NO in the mutant exerts an overriding inhibitory effect on H2-T6SS expression, masking any contributions from lowered cAMP levels. Alternatively, NO accumulation may interfere with the cAMP–Vfr pathway through an as-yet-unidentified mechanism, suggesting the existence of additional regulatory cross-talk between NO signaling and canonical virulence control systems.

Conceptually, this work advances three ideas. First, it identifies a regulatory layer upstream of Dnr, positioning NorCBD as a gatekeeper that sets the usable NO signal for H2-T6SS activation. Second, it supports a biphasic model in which H2-T6SS expression is maximal within an optimal NO window and falls when NO exceeds that range. Third, it links denitrification enzymology to secretion-system regulation and to multiple virulence outputs through changes in second-messenger pools, thereby connecting respiration to contact-dependent antagonism and community behaviors. Importantly, by incorporating recent evidence that Dnr and NorCBD are required for fitness even under normoxia in nitrate-rich environments (Stuut Balsam et al., 2025), our findings broaden the scope of NO homeostasis from a purely anaerobic function to a cross-regime principle that influences *P. aeruginosa* physiology across oxygen gradients.

These findings suggest that modulating NorCBD activity or NO buffering could potentially tune *P. aeruginosa* competitiveness and persistence in both hypoxic and normoxic environments, including CF airways and biofilms. Future studies are required to map the complex genetic interactions among the NO, cAMP–Vfr, and c-di-GMP circuits. Altogether, the data establishes NO homeostasis as a unifying principle for how *P. aeruginosa* integrates redox state with secretion-system regulation and virulence behaviors, with NorCBD acting as an essential balancer that enables Dnr-mediated activation of H2-T6SS across oxygen conditions.

## Data availability statement

### Lead contact

Further information and any requests should be directed to and will be fulfilled by the lead contact, Kangmin Duan.

### Data and code availability

- Data reported in this article will be shared by the lead contact upon request.
- This article does not report the original code.
- Any additional information required to reanalyze the data reported in this article is available from the lead contact upon request.

## Author contributions

M.M.H. and K.D. conceived and designed the study. M.M.H. and S.B. performed the experimentation and generated the data. M.M.H. and K.D. performed data analysis and interpretation. Article writing was performed by M.M.H. and K.D. The final version of the article was approved by all authors.

